# Longitudinal Imaging in a Mouse Model of Focal Cortical Dysplasia Reveals Lesion Expansion and Loss of Pre-Discharge Neuronal Recruitment

**DOI:** 10.64898/2026.10.24.761526

**Authors:** Monika Řehořová, Jana Thao Rozlivková, Jana Populová, Natálie Procházková, Carl Václav Lím Olson, Niki Shariati, Jan Kudláček, Přemysl Jiruška, Ondřej Novák

## Abstract

**Objective:** Focal cortical dysplasia type II (FCD II) is a leading cause of drug-resistant focal epilepsy, yet how its lesional circuits behave and change over time *in vivo* has not been directly observed.

**Methods:** We used a mouse model of FCD II carrying the patient-derived gain-of-function mTOR^p.L2427P^ variant introduced by *in utero* electroporation. Through a chronic cranial window over the lesion, we performed repeated *in vivo* two-photon calcium imaging of identified dysmorphic neurons, pyramidal cells, and VIP interneurons at defined time points, combined with video-EEG monitoring, to determine seizure status and relate cellular activity to interictal discharges.

**Results:** The lesion was not static in adulthood. Dysplastic cortex expanded progressively. This expansion occurred without the tenascin R/C accumulation reported in human FCD II, and was partly explained by continued somatic growth of dysmorphic neurons. Dysmorphic neurons were generally hypoactive; in seizing mice, both dysmorphic and pyramidal cell activity progressively declined, whereas VIP interneuron activity increased slightly in non-seizing animals. In non-seizing lesional cortex, neurons were reliably recruited immediately before interictal discharges. By late disease stages, pre-discharge recruitment had weakened to the point that lesional layer 2/3 neurons were no more active before discharges than expected by chance.

**Significance:** These findings recast FCD II as a lesion that continues to remodel structurally and functionally throughout adulthood, and does so most markedly in animals with seizures. Growth, a cell-type-specific shift in spontaneous activity, and a progressive loss of the lesion’s temporal lead over interictal discharges together suggest that FCD II is a dynamically evolving lesion whose relationship to epileptiform activity changes over time. If the lesion progressively loses its generative role, the window in which removing it can arrest the disease is finite, consistent with clinical evidence favoring earlier surgery.

**Key points:**

- The same neurons inside an FCD II lesion were imaged *in vivo* across epileptogenesis
- Dysplastic cortex expanded over weeks, tending to be larger in seizing mice
- Expansion coincided with dysmorphic neuron growth without bulk tenascin R/C increase
- In seizing mice, dysmorphic and pyramidal neurons became progressively less active
- Pre-discharge neuronal recruitment in non-seizing mice was absent in seizing mice

## INTRODUCTION

Focal cortical dysplasia (FCD) type II is a severe malformation of cortical development and one of the most frequent causes of pharmacoresistant epilepsy in children and young adults. Surgical resection remains the most effective treatment, yet many patients are ineligible or continue to seize post-resection, and no approved mechanism-based therapy exists for FCD II^1, 2^.

FCD II is characterized by cortical dyslamination, dysmorphic neurons (DNs; enlarged, with aberrant neurofilament accumulation) and, in the IIb subtype, balloon cells^1, 3^. Its genetic basis is now largely established to stem from somatic variants that hyperactivate the mTOR pathway, through gain of function in its activators or loss of function in its inhibitors, which produce the malformation^4–7^.

Because the causal variant arises during development, FCD II has been conceptualized largely as a static, developmentally “finished” malformation. This view is difficult to reconcile with the biology of the pathway; mTOR is the canonical regulator of cell size, coupling growth-factor and nutrient signaling to protein synthesis and mass accumulation^8^. DNs are enlarged and show aberrant dendritic arbors and axonal caliber^9, 10^, and human lesional tissue is further remodeled at the level of the extracellular matrix^11^. Whether the dysplastic cortex continues to change structurally in adulthood, and whether such change tracks the emergence of seizures, has not been observed directly.

Much less is known about how the lesion behaves functionally. At the single-cell level, DNs are intrinsically hypoexcitable, requiring more current to reach threshold and firing at reduced frequency^9, 12^, yet they are not electrically inert. Enlarged Ca²⁺ conductances can drive paroxysmal responses once the network recruits them, so low tonic firing does not reflect a fundamental inability to participate in epileptiform activity^13^. How these neurons behave in the intact brain, and whether this changes over time, remains unknown.

In studies sampling a single time point, intrinsic epileptogenicity in FCD II co-localizes with the dysplastic tissue, while interictal abnormalities in surrounding cortex recede once the lesion is removed^14^. Cross-sectional data cannot determine whether the lesion remains the sole generator or progressively recruits other cortical regions, often referred to as secondary epileptogenesis^15^. Classically, a primary focus first drives discharges in connected regions that depend on it. These can mature into independent foci that generate seizures autonomously, and once independent, removing the primary focus no longer stops the seizures^16, 17^. In FCD specifically, longer-standing epilepsy and later surgery are associated with a lower chance of postoperative seizure freedom^18, 19^. If the dysplastic focus behaves likewise—generating pathological activity in the early phase and losing that role as the network reorganizes—the window in which removing the lesion can arrest the disease would be finite.

Here we followed the same lesional neurons over weeks *in vivo*, which cannot be achieved with slice electrophysiology or histology. In a mouse model carrying the patient-derived mTOR^p.L2427P^ variant^4, 10, 20^, we combined chronic two-photon calcium imaging^21, 22^ with continuous video-EEG. We tracked DNs, pyramidal cells (PCs), and vasoactive intestinal peptide–expressing (VIP) interneurons, a disinhibitory population implicated in seizures^23–26^. These recordings show FCD II to be a dynamic lesion, one that continues to remodel in adulthood, structurally and functionally, while its neurons gradually lose their lead over interictal discharges.

## MATERIALS AND METHODS

### Sample preparation

#### Animals and experimental groups

All experiments were performed under the Animal Care and Animal Protection Law of the Czech Republic, in line with EU directive 2010/63/EU, and were approved by the Ethics Committee of the Second Faculty of Medicine (Project License No. MSMT-31765/2019–4). Both male and female mice were included.

The longitudinal cohort comprised 19 VIP-IRES-Cre × Ai9 (LSL-tdTomato) mice, in which tdTomato labels VIP interneurons (Fig. 1C): 14 were electroporated with mutant mTOR (p.L2427P; plasmids from SoVarGen Co., Ltd.^4^, further adapted) and five with wild-type mTOR (WT). Of the mutant mice, eight remained seizure-free throughout video-EEG monitoring (non-seizing, NS), four had their first spontaneous seizure between the early and late imaging windows (seizure-onset, SO), and two were already seizing when imaging began (already-seizing, AS). A separate histological cohort of five mutant mice (C57BL/6J background), all with behaviorally confirmed seizures (see *Supplementary Methods*), was used only for tenascin immunohistochemistry.

**Figure 1.**
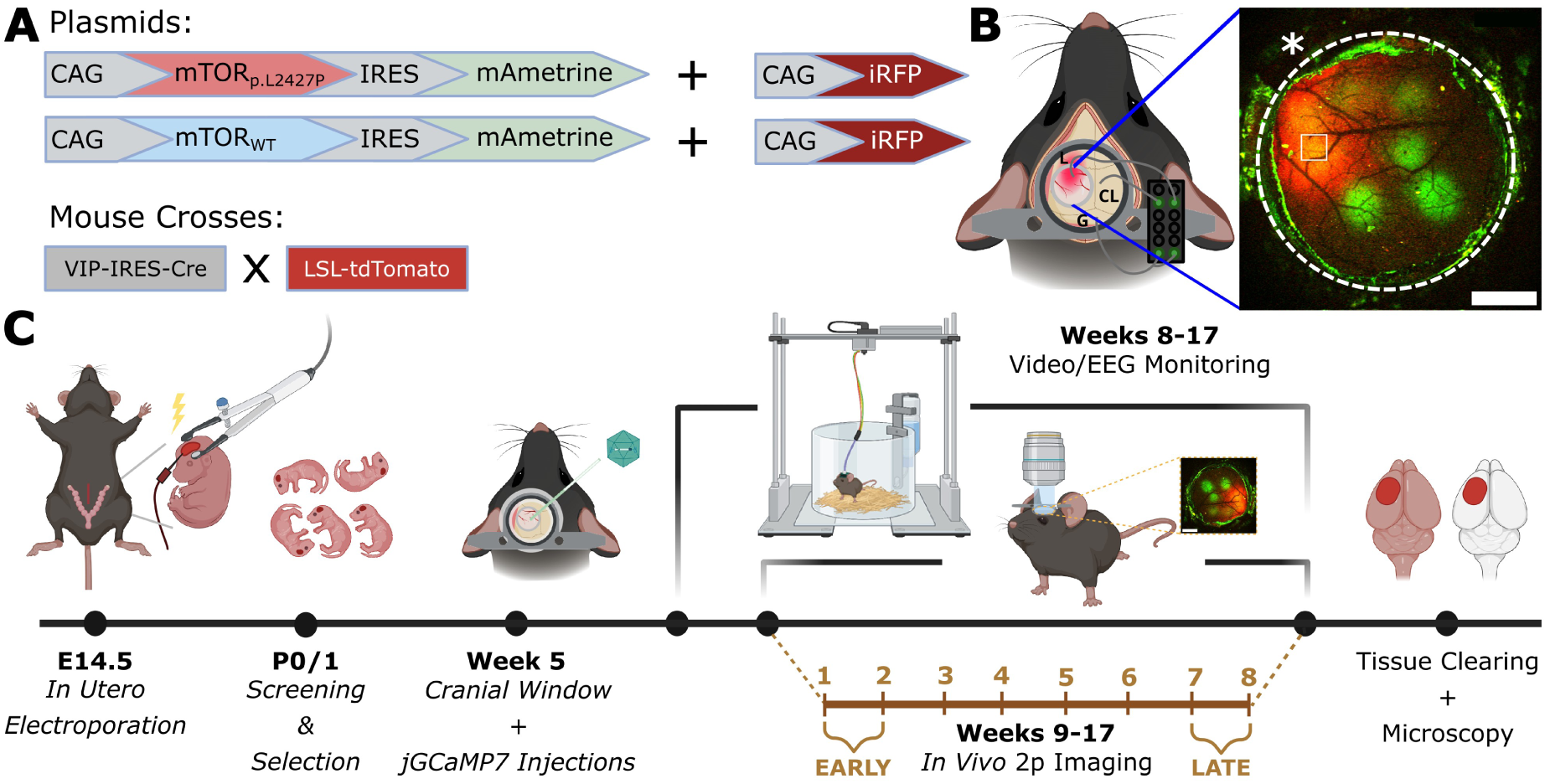
Experimental design and timeline. (**A**) Constructs and mouse crosses. FCD was induced by *in utero* electroporation of a CAG-driven cassette co-expressing mutant mTOR (mTOR^p.L2427P^; wild-type mTOR in controls) with the mAmetrine reporter, together with a separate CAG-iRFP plasmid that provided a near-infrared label for transcranial localization of the electroporated area. VIP-IRES-Cre mice were crossed with LSL-tdTomato reporters to drive tdTomato selectively in VIP interneurons. (**B**) Dorsal schematic of the cranial window over the FCD lesion (carmine). The titanium headbar carries a right-sided connector to which three platinum electrodes are attached: two LFP electrodes, one in the lesion (L), and one in the contralateral hemisphere (CL), plus a ground (G). Inset: representative macroscopic fluorescence image showing the spatial overlap of the iRFP-labeled electroporated area (red) and the jGCaMP7f injection sites (green). The dashed circle marks the 4 mm cranial window, the square a 405 × 405 µm imaging field of view (FOV), and the asterisk the typical position of the lesional electrode relative to the window and the imaged FOVs. Scale bar = 1 mm. (**C**) Experimental pipeline: *in utero* electroporation at E14.5; screening and selection of electroporated pups at P0/1; cranial window implantation with jGCaMP7f AAV injection at week 5; longitudinal *in vivo* two-photon imaging (weeks 9–16) paired with continuous video-EEG monitoring (weeks 8–16); and terminal tissue clearing with light-sheet microscopy. Weekly imaging sessions are numbered 1– 8 (relative to imaging onset) and grouped into EARLY (imaging sessions 1+2) and LATE (imaging sessions 7+8) windows for longitudinal comparison. Panels B and C were created with BioRender.com.

#### Surgery and tissue processing

*In utero* electroporation (IUE) was performed at embryonic day 14.5 as previously described^10^ with a custom-built electroporator^27^, injecting pCAG-iRFP_713_ with either pCAG-mTOR_WT_ or pCAG-mTOR_MUT_ into the left lateral ventricle (pCAG-EGFP replaced pCAG-iRFP_713_ in the histological cohort); only pups with a clearly unilateral lesion in the somatosensory cortex were retained. At postnatal weeks 5–6, a titanium headbar with an electrode connector was attached, a 4.5 mm craniotomy was made over the lesion, AAV9-syn-jGCaMP7f was injected at 10 sites in layer 2/3 within and outside the lesion, and the craniotomy was sealed with a 4 mm coverslip on the intact dura. At 17 weeks, mice were perfused with 4% paraformaldehyde. Coronal sections of the histological cohort were immunostained for tenascin-R or tenascin-C^10^, and six brains of the functional cohort (two WT, four mutant) were cleared with CUBIC^28^. Details are given in *Supplementary Methods*.

### Data acquisition

#### Video-EEG monitoring

From postnatal week 8–9 until week 16, mice underwent continuous video-EEG monitoring, interrupted only by the weekly imaging sessions. All analyzed EEG signals came from an electrode in the FCD lesion, 1–3 mm from the imaged fields, with a cerebellar reference/ground. Video was used only to verify electrographic seizures.

#### Imaging

From one week after the start of monitoring, mice were imaged weekly under awake head fixation with simultaneous EEG, on a custom-built two-photon microscope (920 nm; 20×/1.0 NA objective; ScanImage interface^29^). In each mouse, 6–10 sites were selected on a low-magnification overview showing the iRFP-labeled lesion and relocated in every session. Each field of view (FOV; 405 × 405 µm, 150–300 µm below the dura) was recorded twice for 5 min per session at 30 frames/s and classified relative to the iRFP-delineated lesion border as lesional (fully inside), edge, or extra-lesional (“out”). Imaging and EEG were synchronized within one frame (33 ms).

Immunostained sections were imaged on a spinning-disk confocal microscope with identical settings for both hemispheres (two sections per antibody per animal). Cleared brains were imaged by light-sheet microscopy for three-dimensional visualization and a descriptive count of VIP interneuron density.

### Data processing and statistics

#### Imaging windows

Of the eight imaging sessions, EARLY denotes sessions 1–2 and LATE sessions 7–8, about six weeks apart; functional measures were averaged across the two sessions of each window. Cortical expansion and soma size were measured between sessions 2 and 8. In SO animals, the EARLY window preceded and the LATE window followed the first seizure. AS animals contribute to the expansion and soma-size comparisons (group S, with SO), to the time-since-seizure analysis (Supplementary Fig. 3), and to the late peri-IED data, but not to the EARLY-versus-LATE comparisons of firing.

#### Calcium signal processing

Image sequences were motion-corrected with NoRMCorre^30^ and regions of interest (ROIs) from a reference session were mapped to all others. VIP interneurons were identified by tdTomato; the remaining neurons were classified by soma size as DNs (>400 pixels, ∼18 µm) or PCs (<300 pixels, ∼15 µm); intermediate ROIs were excluded. Spiking was inferred from neuropil-corrected ΔF/F traces with CASCADE^31^ (GCaMP8-tuned model^32^); traces with noise ≥5 %·Hz^−1/2^ were discarded. The mean inferred rate per neuron is termed the spontaneous firing rate (SFR).

#### Interictal discharges

IEDs were detected in the local field potential (LFP) recorded during imaging^33^, verified manually, and timed at their peak deflection. Inferred spiking was summed over the five frames (∼167 ms) before (pre-) and after (post-) each IED, excluding the discharge frame. Per FOV (≥3 qualifying neurons), the fraction of neurons with ≥1 discrete CASCADE spike in the five pre-IED frames was compared with that of the same neurons at an equal number of random times ≥60 frames (2 s) from any IED.

#### Structural measures

Expansion along the mediolateral (ML) and anteroposterior (AP) axes was obtained from affine registration (itk-elastix^34, 35^) of averaged FOV images from sessions 2 and 8. Soma size was measured in five FCD mice on 60 µm Z-stacks from sessions 2 and 8. An annotator blinded to session order delineated the 3–4 largest (presumed DN) and 3–4 smallest (presumed PC) neurons present in both stacks, and the effective diameter was derived from the largest cross-sectional area. Tenascin was quantified on maximum-intensity projections as the mean-intensity ratio between the EGFP-defined lesion and a homotopic contralateral region, delineated blind to the tenascin channel, and averaged to one value per animal.

#### Statistical analyses

Unless stated otherwise, the unit of analysis is the FOV, with one value per condition (imaging window, group, or time-since-seizure bin) averaged across neurons of a given type and across recordings. Firing rates were analyzed with linear mixed-effects models with random intercepts for animal and, except for the cross-sectional VIP comparison, FOV (MATLAB R2024a, fitlme, restricted maximum likelihood), with Satterthwaite degrees of freedom for contrasts and likelihood-ratio tests for omnibus effects. Other comparisons used one-sample, paired, or Welch’s *t*-tests and, for active-neuron fractions, permutation tests (10^5^ resamples). All tests were two-sided and families of related comparisons were Holm-corrected. Values are means or model-estimated marginal means ± SEM. Model specifications, test families, and sample sizes are in *Supplementary Methods*. Significance: \**p* < 0.05, \*\**p* < 0.01, \*\*\**p* < 0.001; ns, not significant.

## RESULTS

We used a previously established model of FCD II generated by IUE of the patient-derived mTOR^p.L2427P^ gain-of-function variant or of wild-type mTOR in controls (Fig. 1A). A co-electroporated near-infrared reporter (iRFP_713_) delineated the electroporated area through the cranial window, leaving the green channel for jGCaMP7f and the red channel for tdTomato-labeled VIP interneurons. Each mouse carried a lesional and a contralateral LFP electrode for continuous video-EEG; the lesional electrode was placed at the edge of the 4 mm cranial window, 1–3 mm from the imaged fields (Fig. 1B, asterisk and square), and the same FOVs were imaged weekly over eight sessions with simultaneous LFP (Fig. 1C). Mutant mice were grouped by seizure history as non-seizing (NS), seizure-onset (SO; first seizure between the early and late imaging windows), or already-seizing (AS); the epileptic phenotype of the seizing animals (2 AS, 4 SO; 4.8 ± 2.9 seizures per day, mean duration 49.0 ± 3.4 s) is described in Supplementary Fig. 1.

### Lesional cortex expands progressively in seizing mTOR-mutant mice

To ask whether FCD lesions keep remodeling in adulthood, we registered each FOV imaged at session 2 onto the same FOV at session 8 and read the tissue deformation from the fitted affine transformation (Fig. 2A, B). In WT cortex, the two time points were essentially superimposable, whereas lesional FOVs of seizing mice (S = SO + AS) were radially enlarged. S FOVs expanded by 3.0 ± 0.5% mediolaterally (ML) and 2.7 ± 0.5% anteroposteriorly (AP; Holm-adjusted *p* < 0.001 versus 0%), whereas neither NS (ML: 1.0 ± 0.5%; AP: 1.3 ± 0.5%) nor WT FOVs (ML: −0.3 ± 0.7%; AP: −1.5 ± 0.6%) differed from zero (all adjusted *p* ≥ 0.13; Fig. 2C). S exceeded WT on both axes (ML: adjusted *p* = 0.022, Hedges’ *g* = 1.39; AP: adjusted *p* = 0.001, *g* = 1.94) and NS exceeded WT anteroposteriorly only (adjusted *p* = 0.023, *g* = 1.19; ML: adjusted *p* = 0.33); S and NS did not differ significantly on either axis (ML: adjusted *p* = 0.051, *g* = 0.83; AP: adjusted *p* = 0.26).

**Figure 2.**
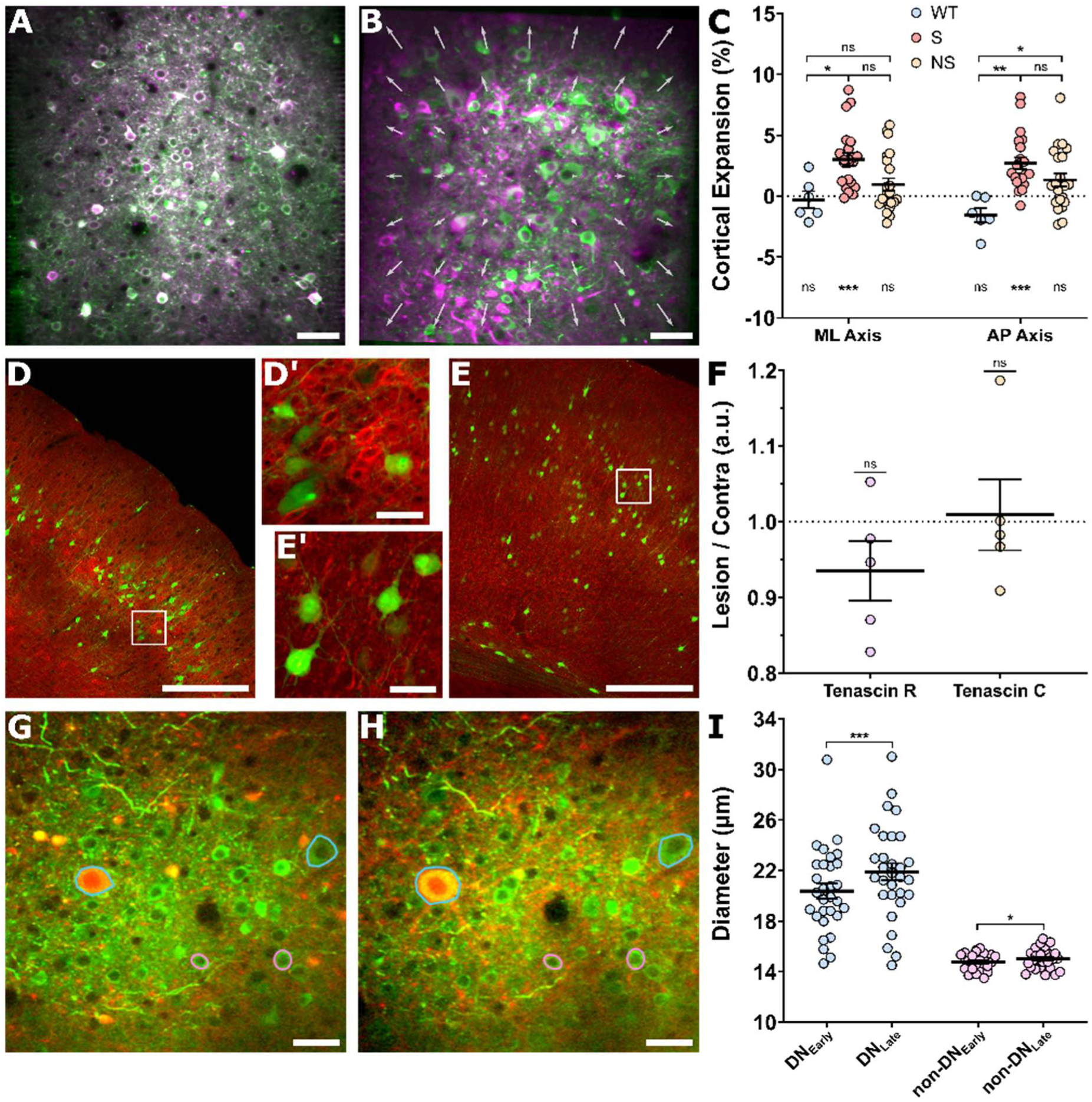
Progressive, anisotropic cortical expansion is specific to seizing mTOR-mutant mice and accompanies growth of DNs. (**A**, **B**) Overlays of the same FOV imaged at session 2 (green) and session 8 (magenta) in a mTOR_WT_ mouse (**A**) and a seizing mTOR_MUT_ mouse (**B**). White arrows in (**B**) indicate the direction and magnitude of the fitted affine transformation aligning the two timepoints, revealing expansion of the lesional tissue that is absent in the WT control. (**C**) Cortical expansion (%) relative to the baseline session, measured along the mediolateral (ML) and anteroposterior (AP) axes over six weeks in adulthood, for wild-type controls (WT, blue), mTOR-mutant mice that developed spontaneous seizures (S, pink), and mTOR-mutant mice without seizures (NS, orange). Horizontal bars, group means. Brackets denote between-group comparisons; markers below each cluster test each group against no change (vs. 0%). (**D**, **E**) Confocal imaging and max-intensity projections of the lesion immunolabeled for two extracellular-matrix proteins, tenascin R (**D**) and tenascin C (**E**). Green, (predominantly) DNs within the lesion; red, CF647-conjugated antibody against the respective tenascin. Higher-magnification insets (**D′**, **E′**) show no prominent association between tenascin labeling and the green neurons. (**F**) Bulk tenascin-antibody fluorescence within the FCD lesion expressed as a ratio to the homotopic contralateral cortex (Lesion/Contra, a.u.). Neither tenascin R nor tenascin C differed significantly from the contralateral side. Each point is one animal. (**G**, **H**) Representative two-photon FOV showing the morphological and size changes of individual neurons between an early (**G**; session 2) and a late (**H**; session 8) imaging session. Two DNs are outlined in cyan and two normal-sized PCs in pink. (**I**) Effective soma diameter of DNs and normal PCs at the early versus late imaging time points; horizontal bars, means. Scale bars: **A**, **B** = 100 μm; **D′**, **E′**, **G**, **H** = 50 μm; **D**, **E** = 500 μm. Bars represent arithmetic means, whiskers SEM.

### Expansion is not accompanied by tenascin R or tenascin C accumulation

Human FCD type II is characterized by a marked accumulation of tenascin R and tenascin C in the lesional neuropil^11^. In a separate cohort of five seizing FCD mice, in which electroporated neurons were labeled with EGFP (*Methods*), tenascin immunoreactivity showed no prominent association with the EGFP⁺ neurons (Fig. 2D, E and insets D′, E′), and the lesion-to-contralateral intensity ratio differed from unity for neither tenascin R (0.94 ± 0.04; *t*(4) = −1.64, *p* = 0.35) nor tenascin C (1.01 ± 0.05; *t*(4) = 0.20, *p* = 0.85; Fig. 2F; positive controls in Supplementary Fig. 2). Unlike in human FCD II, the expansion is thus not accompanied by a detectable bulk accumulation of either tenascin.

### Dysmorphic neurons continue to grow in adulthood and account for part of the expansion

We next asked whether continued growth of the mutant neurons could contribute to the aforementioned cortical expansion. Tracking the same somata across the imaging period (Figs. 2G, H), DNs enlarged from an FOV-mean diameter of 20.4 ± 0.6 µm to 21.9 ± 0.7 µm (Holm-corrected *p* < 0.001; increase seen in 23 of 30 FOVs), whereas PCs changed only marginally (14.7 ± 0.1 µm to 15.0 ± 0.2 µm; *p* = 0.015; 20 of 26 FOVs; Fig. 2I). Within fields containing both classes, DNs grew 1.49 ± 0.38 µm more than PCs (*p* < 0.001; positive in 20 of 26 fields). Ongoing enlargement of the mutant neurons thus contributes to the expansion, although it is unlikely to account for its full magnitude (see *Discussion*).

### Cell type and seizure history shape how spontaneous firing is remodeled in and around the lesion

We then quantified the spontaneous firing rate (SFR) of individual neurons, inferred from ΔF/F traces (illustrative image in Fig. 3A), in DNs, PCs, and VIP interneurons identified within each field (Fig. 3B; representative traces in Fig. 3C). At baseline, DNs were markedly hypoactive. Within the lesion, their firing rate was about one-third lower than that of neighboring PCs (estimated marginal means 0.54 ± 0.06 vs. 0.79 ± 0.06 Hz; Holm-corrected *p* < 0.001) and also lower than that of PCs outside the lesion (0.78 ± 0.06 Hz; *p* < 0.001), whereas PCs fired at the same rate inside and outside the lesion (*p* = 0.87; Fig. 3D). VIP firing did not differ between the lesion and surrounding cortex (model means 0.58 ± 0.10 vs. 0.48 ± 0.10 Hz; *p* = 0.25; Fig. 3E). The apparent gap in raw FOV averages (0.45 vs. 0.60 Hz) reflected the uneven distribution of VIP FOVs across mice.

**Figure 3.**
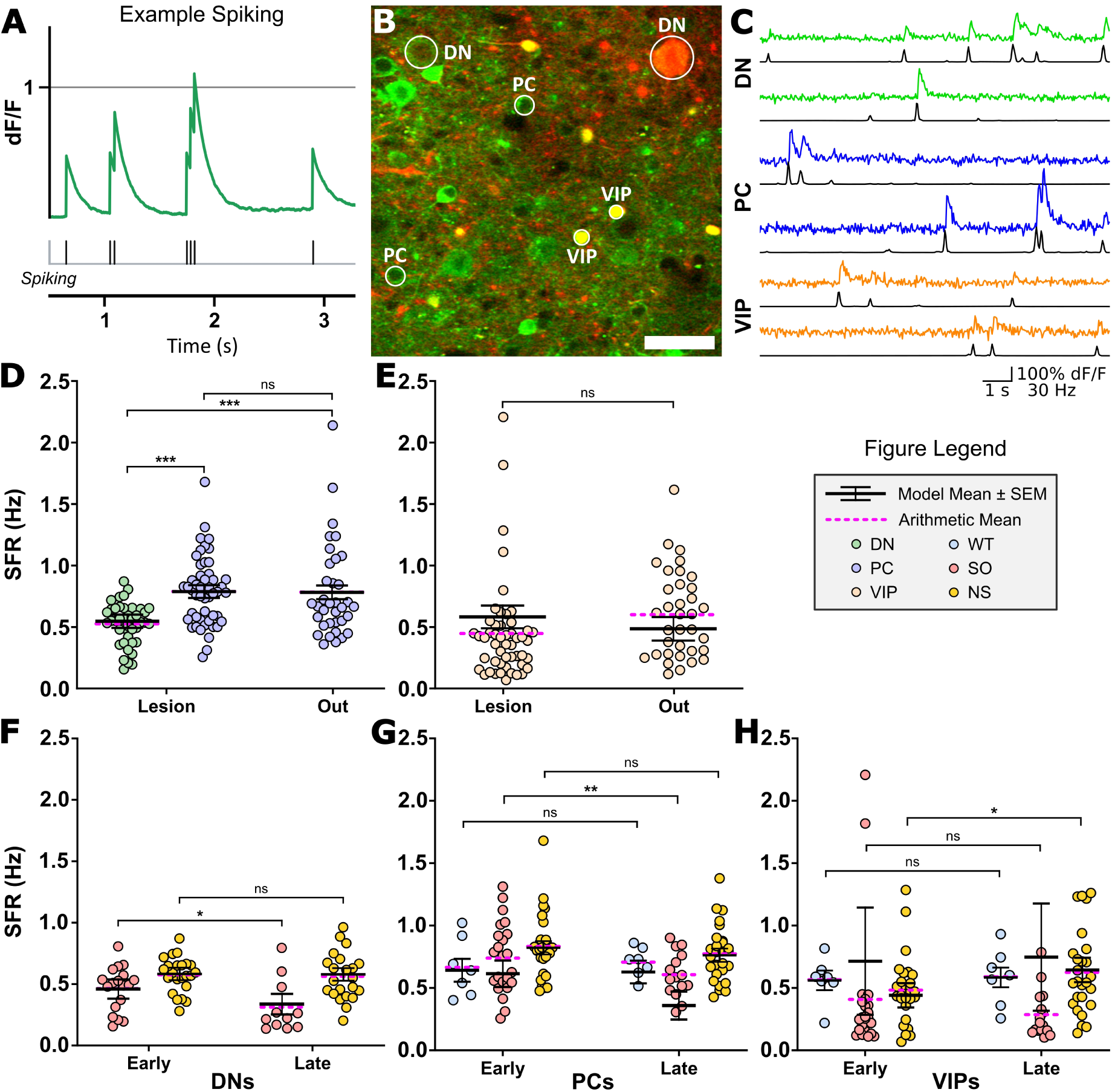
Cell-type- and seizure-history-specific changes in spontaneous firing in and around the lesion. (**A**) Principle of spike inference from the calcium signal; an idealized ΔF/F trace (top) and the spike train inferred from it (bottom; each tick is one inferred spike, shown here in binary form). (**B**) Representative two-photon image of a lesional field, cropped to 250 × 250 µm; green, jGCaMP7f; red, iRFP_713_ (recorded in the tdTomato emission channel); yellow, VIP interneurons co-expressing jGCaMP7f and tdTomato. Circles mark examples of the analyzed cell types: dysmorphic neurons (DN), pyramidal cells (PC) and VIP interneurons (VIP). Scale bar, 50 µm. (**C**) Representative ΔF/F traces of DNs (green), PCs (blue), and VIP interneurons (orange), each with the CASCADE-inferred firing rate beneath (black). Scale bars = 1 s, 100% ΔF/F, and 30 Hz. (**D**–**H**) Spontaneous firing rate (SFR, Hz), one value per FOV per cell type (the mean of its constituent neurons). Solid horizontal bars, model-estimated marginal means from linear mixed-effects models with random intercepts for mouse and, except in E, FOV (see *Supplementary Methods*); whiskers, SEM; dashed magenta bars, arithmetic FOV means. Brackets denote Holm-corrected pairwise contrasts (ns, not significant). (**D**) DNs (green) and PCs (lavender) in lesional versus extra-lesional (“out”) FOVs; SO and NS mice pooled, early window (DN-lesion n = 41, PC-lesion n = 50, PC-out n = 35 FOVs; 12 mice). (**E**) VIP interneurons in lesional versus extra-lesional FOVs (SO and NS pooled, early window; lesion n = 50, out n = 35 FOVs; 12 mice); model and arithmetic means diverge because FOVs were unevenly sampled across mice (see *Results*). (**F**–**H**) SFR in lesional FOVs at the early versus late imaging window, split by seizure-history group (WT, light blue; SO, red; NS, yellow), for (**F**) DNs, (**G**) PCs, and (**H**) VIP interneurons. Brackets denote the early-versus-late contrast within each group, Holm-corrected across groups within the panel. The SO VIP estimate in (**H**) is not interpreted: with four animals its standard error (0.44 Hz) is four- to fivefold larger than in the WT and NS groups (0.09 and 0.11 Hz). FOV numbers (early/late): **F**, SO 18/11 and NS 23/22; **G**, WT 7/7, SO 23/15 and NS 27/27; **H**, WT 7/7, SO 23/16 and NS 27/26; 3 WT, 4 SO and 6 NS mice. In WT animals, which develop no lesion, FOVs lay within the iRFP-labeled electroporated area.

Within the lesion, firing in WT controls was stable over the six weeks for both PCs (0.65 ± 0.10 → 0.69 ± 0.10 Hz; *p* = 0.56) and VIP interneurons (0.56 ± 0.09 → 0.58 ± 0.09 Hz; *p* = 0.68), indicating no systematic drift from the preparation or repeated imaging. In mutant animals, excitatory neurons changed only in SO mice. DNs, already the least active population, decreased their activity further (0.51 ± 0.09 → 0.35 ± 0.09 Hz; *p* = 0.014; NS: 0.60 ± 0.06 → 0.60 ± 0.06 Hz, *p* = 0.93; Fig. 3F), and PC firing fell markedly (0.69 ± 0.12 → 0.44 ± 0.12 Hz; *p* = 0.003; NS: 0.84 ± 0.06 → 0.78 ± 0.06 Hz, *p* = 0.46; Fig. 3G). VIP interneurons changed in the opposite direction and only in NS mice, where their firing rate increased (0.51 ± 0.11 → 0.65 ± 0.11 Hz; *p* = 0.044). VIP firing was unchanged in WT mice, and the SO estimate was too unstable to interpret (Fig. 3H). The late phase in SO mice was therefore marked by suppression of excitatory neuronal activity, whereas the only late increase in firing occurred in VIP interneurons of non-seizing mice. This suppression was not a post-ictal effect. In seizing animals, pyramidal firing within 5 hours of a seizure was only 0.078 Hz (95% CI 0.020–0.14) lower than at longer intervals, about a third of the 0.25 Hz early-to-late decline, and dysmorphic and VIP firing did not depend on seizure proximity (Supplementary Fig. 3).

VIP interneuron density, counted in cleared whole brains, was approximately 45% lower in mutant than in wild-type cortex (970 ± 160 vs 1770 ± 50 neurons/mm^3^; n = 4 mutant and 2 WT brains; Supplementary Fig. 4). The WT value is comparable to published estimates for mouse somatosensory cortex^36^. Given the expanded cortex, the two control brains and the reduction in overall neuronal density previously reported in this model^10^, we report these values descriptively and do not read them as a loss specific to VIP interneurons.

### Lesional neurons are recruited before interictal discharges and this recruitment declines across epileptogenesis

Imaging lesional fields allowed us to relate single-cell firing to individual interictal discharges (IEDs). Subtracting a pre-IED baseline image from the average of frames around the discharge showed that IEDs were accompanied by a diffuse neuropil signal on which the brightening of individual somata could be resolved (Fig. 4A–C). In an example NS field, DNs, PCs, and VIP interneurons showed transient increases in firing that clustered around the LFP-detected IED and rose just before its peak (Fig. 4D). Across fields of non-seizing animals, activity in the five frames preceding the IED exceeded that in the five frames following it in all three classes (Fig. 4E; Holm-corrected within condition: DN 0.72 ± 0.16 → 0.19 ± 0.04 a.u., *p* = 0.004; PC 0.60 ± 0.11 → 0.21 ± 0.02, *p* = 0.003; VIP 0.36 ± 0.07 → 0.18 ± 0.03, *p* = 0.01). Early-phase tissue of SO animals showed the same direction of effect, which did not survive statistical correction (DN *p* = 0.16, PC *p* = 0.16, VIP *p* = 0.17). In late-stage tissue, the lead was lost. DNs showed no pre–post difference (*p* = 0.97), VIP activity was higher after the discharge (*p* = 0.02), and PCs trended in the same direction (*p* = 0.07). Extra-lesional fields showed no pre-IED structure in any condition (all *p* ≥ 0.09), so IED-leading activity was specific to the lesion.

**Figure 4.**
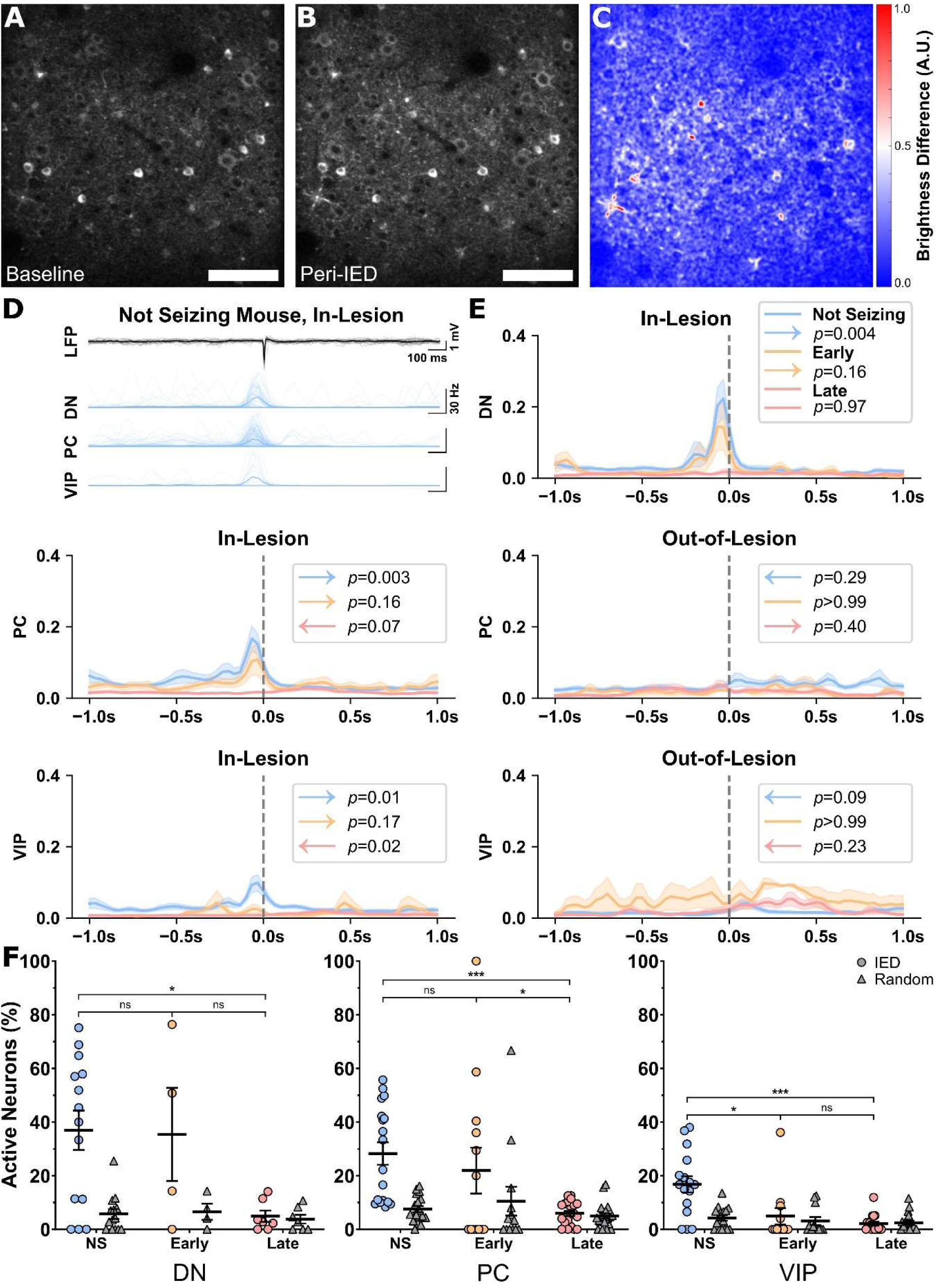
Lesional neurons lead IEDs in non-seizing tissue and this lead is lost across epileptogenesis. (**A**–**C**) Example in-lesion field during an interictal discharge. (**A**) Baseline image, averaged over 20–30 frames before the IED. (**B**) Peri-IED image, averaged over frames −5 to +5 around the IED. (**C**) Difference between (**B**) and (**A**), normalized to 0–1 (a.u.) for display. Scale bars = 100 µm. (**D**) Example non-seizing, in-lesion field. Top, overlaid LFP traces centered on manually verified IEDs (scale bars = 100 ms, 1 mV); bottom, CASCADE-inferred instantaneous firing rate of individual DNs, PCs, and VIP interneurons aligned to the IED peak (scale bar = 30 Hz). (**E**) Mean ± SEM peri-IED firing rate (a.u.) aligned to the IED at t = 0 (dashed line), in-lesion (DN, PC, VIP) and extra-lesional (PC and VIP only). Color denotes condition: non-seizing (blue), early (orange), and late (pink) seizing tissue. Arrows indicate whether integrated activity was higher before (→) or after (←) the IED. *P-*values are from two-sided one-sample *t*-tests of the per-FOV pre-minus-post difference against zero, Holm-corrected across cell types within each condition (see *Methods*). (**F**) Fraction of active neurons per field (%), defined as neurons with ≥1 CASCADE-inferred spike in the five frames preceding an event (discrete CASCADE output). Each symbol is one FOV: colored circles, real IEDs; grey triangles, the same neurons at random times; horizontal bars, across-FOV means ± SEM. Symbols are grouped by cell type and by condition, color-coded as in (**E**). Brackets compare the real-IED active fraction between conditions; *p-*values are from two-sided permutation tests (label shuffling, 10⁵ resamples), Holm-corrected across the three pairwise comparisons within each cell type. Random-time controls are tested in Supplementary Fig. 5.

To distinguish weaker per-neuron responses from fewer participating neurons, we computed per FOV the fraction of neurons with ≥1 inferred spike in the 167 ms (five frames) before each IED and in the same neurons at random times (Fig. 4F). In non-seizing tissue, more neurons were active before real IEDs than at random times in all three classes (DN *p* = 0.003, PC *p* < 0.001, VIP *p* < 0.001; Supplementary Fig. 5). This excess was absent in early and late tissue (all *p* > 0.99). The real-IED active fraction declined from non-seizing to late tissue in PCs and VIPs (both *p* < 0.001) and in DNs (*p* = 0.02). In VIPs, it was reduced already in the early phase (*p* = 0.02) and in PCs an early-to-late reduction was observed at borderline significance (21.9 ± 8.6% → 5.9 ± 0.9%; *p* = 0.049). Interpreted cautiously, these data suggest that the pre-IED lead is lost because fewer lesional neurons are recruited around the discharge, rather than because individual neurons respond more weakly.

## DISCUSSION

Using a chronic cranial window in a mouse model of FCD II based on a patient-derived variant, we followed identified neurons inside the lesion across weeks of epileptogenesis with concurrent video-EEG. Three linked findings emerged: the FCD lesion continues to remodel structurally in adulthood, its neurons restructure their firing in a manner specific to cell type and seizure history, and the lesion progressively uncouples from the interictal activity it is thought to generate.

### A lesion that continues to grow

FCD type II is classically framed as a static, developmentally “finished” malformation^1, 2^. The progressive expansion of dysplastic cortex in adult mice argues instead for an ongoing process. Expansion was most pronounced in animals that developed seizures, but the association may reflect lesion severity rather than seizures themselves. More severe lesions carry a higher seizure risk and a higher load of DNs may themselves drive remodeling. Extra-lesional fields, which could separate the two, were too few in seizing animals. The most parsimonious cellular contributor is continued growth of the DNs, which enlarged while neighboring PCs remained nearly static. This fits the core biology of the pathway—mTOR as the canonical regulator of cell size^8^—and reframes adult somatic growth as a candidate substrate for disease progression, rather than a purely developmental feature. Somatic growth is, however, unlikely to explain the full deformation. Unlike human FCD II^11^, the model showed no bulk tenascin accumulation. The expansion may also involve other processes that we did not measure, such as reactive astrogliosis^11, 37^, neuritic growth^10^, and other extracellular-matrix constituents^11^.

### Firing is remodeled according to cell type and seizure history

At baseline, DNs were markedly hypoactive relative to neighboring PCs. This agrees with human FCD electrophysiology, in which DNs show reduced functional integration, raising the possibility that they are not the direct pacemakers of ictogenesis^13, 38^ and slice data showing that DNs require more current to reach threshold^9, 12^. Why, then, is the lesion epileptogenic? A low mean rate constrains only the average, not how these cells reach threshold or when they fire. DNs carry larger Ca^2+^ current densities^13^ and they may operate in an unstable Cl^-^environment in which synchronous GABAergic drive becomes excitatory^9^; such drive sustains interictal discharges in human FCD tissue^39^. Consistent with this, network stimulation recruits DNs at a lower threshold than their neighbors^12^. Our data extend this dissociation to the intact cortex. The tonic deficit persists in the intact network, yet the same neurons are readily recruited by interictal discharges. Low spontaneous firing should therefore not be read as low pathological relevance.

In seizing animals, both dysmorphic and pyramidal populations became progressively less active. This was not a post-ictal state. Recordings close to a seizure showed only modestly lower pyramidal firing, about a third of the longitudinal decline, and dysmorphic firing was unaffected by seizure timing. One plausible interpretation is homeostatic. Cortical networks defend activity set-points against chronic perturbation^40^, and hyperpolarizing mutant neurons reduces seizure burden^41^, so the suppression may be compensatory, yet insufficient, since seizures and interictal discharges persisted. This suppression may also be clinically relevant; FCD lesions are typically hypometabolic on interictal FDG-PET^42^ and reduced neuronal output may contribute to that signature. VIP interneurons, disinhibitory nodes that gate cortical gain^23, 26^, increased their firing only in non-seizing animals, the only mutant group in which neither excitatory population declined. Because PV and SST interneurons were not recorded, we cannot tell whether this change contributes to that stability, follows from it, or merely accompanies it.

### Decoupling from epileptiform dynamics

In non-seizing lesional tissue, all three cell classes were active before interictal discharges. By the chronic phase, this lead was lost and the fraction of neurons recruited fell to that at random times. The residual post-discharge weighting in VIP and PCs is thus a remnant within a largely unresponsive population, rather than a new follower role. We are cautious about equating the loss of a somatic spike lead in layer 2/3 with a loss of generative role. The lesion may continue to drive discharges through deeper layers, through a minority of neurons below our detection threshold, or through GABAergic synaptic mechanisms^39^. Source-level analysis of the discharge will be needed to distinguish a genuine transfer of generative activity to other cortex from these possibilities, and because extra-lesional fields never showed pre-IED structure, our data localize where the lead was lost, not where it was gained.

Clinically, epileptogenicity in FCD often extends beyond the lesion, and the further it extends, the worse the prognosis. In stereo-EEG of 36 patients with FCD or dysembryoplastic tumors, only 31% had a strictly focal organization, 61% network, and 8% bilateral, and seizure freedom followed this gradient at 87%, 57%, and 0%^43^. In surgical series, older age at surgery and longer epilepsy duration predict lower seizure freedom in FCD^18^ and in other malformations of cortical development^44^. Where these studies infer the extent of epileptogenicity from what survives resection, we observe a transition directly, in identified cells, as a loss of temporal precedence. If the lesion progressively cedes its generative role, the therapeutic window is finite. Our data cannot establish that this loss is what closes the window, but they identify a measurable correlate of the transition.

### Limitations and outlook

Our measurements are correlational; we did not manipulate the pathway or circuit, so whether excitatory suppression is protective, maladaptive, or incidental remains open. Calcium-inferred spiking is a filtered view of firing and a single patient-derived variant cannot capture the mosaic heterogeneity of human FCD II. Within these limits, the study indicates that FCD lesions are dynamically evolving structures—growing, altering its excitatory–inhibitory balance, and shifting its temporal relationship to its own epileptiform output—and provides a basis for testing whether mTOR inhibition halts adult growth and arrests the functional decoupling.

## ACKNOWLEDGEMENTS

We thank Dr. Carsen Stringer from the Janelia Research Campus, Howard Hughes Medical Institute, for her consultations and assistance with processing calcium traces into spiking events. We are also grateful to Dr. Monika Žárská for providing the glioma spheroid used for the validation of the tenascin-C antibody. Finally, we thank Jiří Černý and the Light Microscopy Core Facility of the Institute of Molecular Genetics of the Czech Academy of Sciences (LMCF IMG) for their assistance with light-sheet data acquisition. LMCF IMG is supported by the Ministry of Education, Youth and Sports of the Czech Republic (projects LM2023050, CZ.02.1.01/0.0/0.0/18_046/0016045, and CZ.02.01.01/00/23_015/0008205 – Czech-BioImaging).

## AUTHOR CONTRIBUTIONS

According to CRediT taxonomy the contributions of individual authors were as follows: Conceptualization: MŘ, JTR, and ON. Methodology: MŘ, JTR, NP, CVLO, JK, and ON. Software: JTR, MŘ, JK, and ON. Formal analysis: MŘ, JTR, JP, NP, CVLO, NS, JK, and ON. Investigation: MŘ, JTR, JP, NP, CVLO, NS, JK, and ON. Data curation: MŘ, JTR, JP, CVLO, JK and ON. Writing: MŘ, JTR, JP, NP, JK, and ON. Visualization: MŘ, JTR, CVLO, and ON. Supervision: PJ and ON. Project administration: MŘ, PJ, and ON. Funding acquisition: MŘ, JP, NP, JK, PJ, and ON.

## DATA AVAILABILITY

The majority of the original raw data will be deposited in Zenodo upon publication. The large calcium imaging dataset (>100 TB; motion-corrected only) is available upon reasonable request from the corresponding author.

## ETHICS STATEMENT

All experiments were performed under the Animal Care and Animal Protection Law of the Czech Republic, in line with EU directive 2010/63/EU, and were approved by the Ethics Committee of the Second Faculty of Medicine (Project License No. MSMT-31765/2019–4).

## SUPPLEMENTARY MATERIAL

**Supplementary Figure 1.**
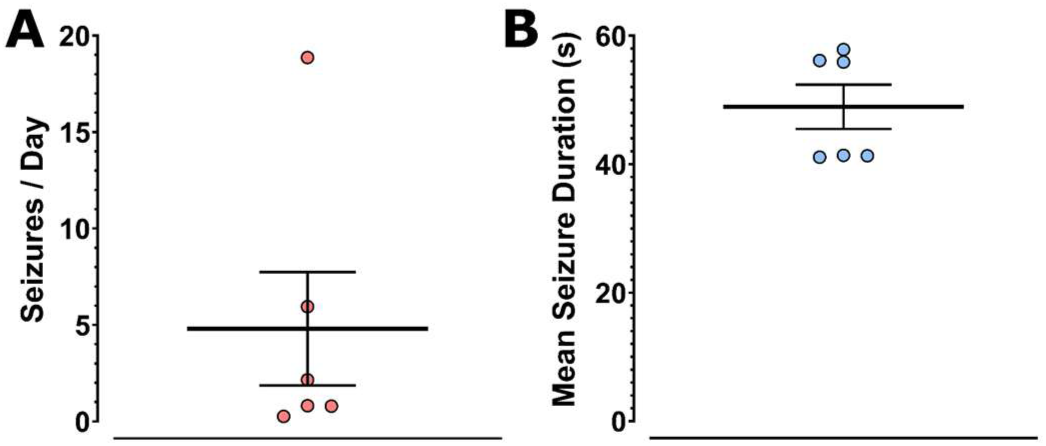
Animals with seizures involved in the study. **(A)** Number of seizures per day and their mean duration **(B)** for all seizing mice included in this project; points denote individual mice; mean ± SEM.

**Supplementary Figure 2.**
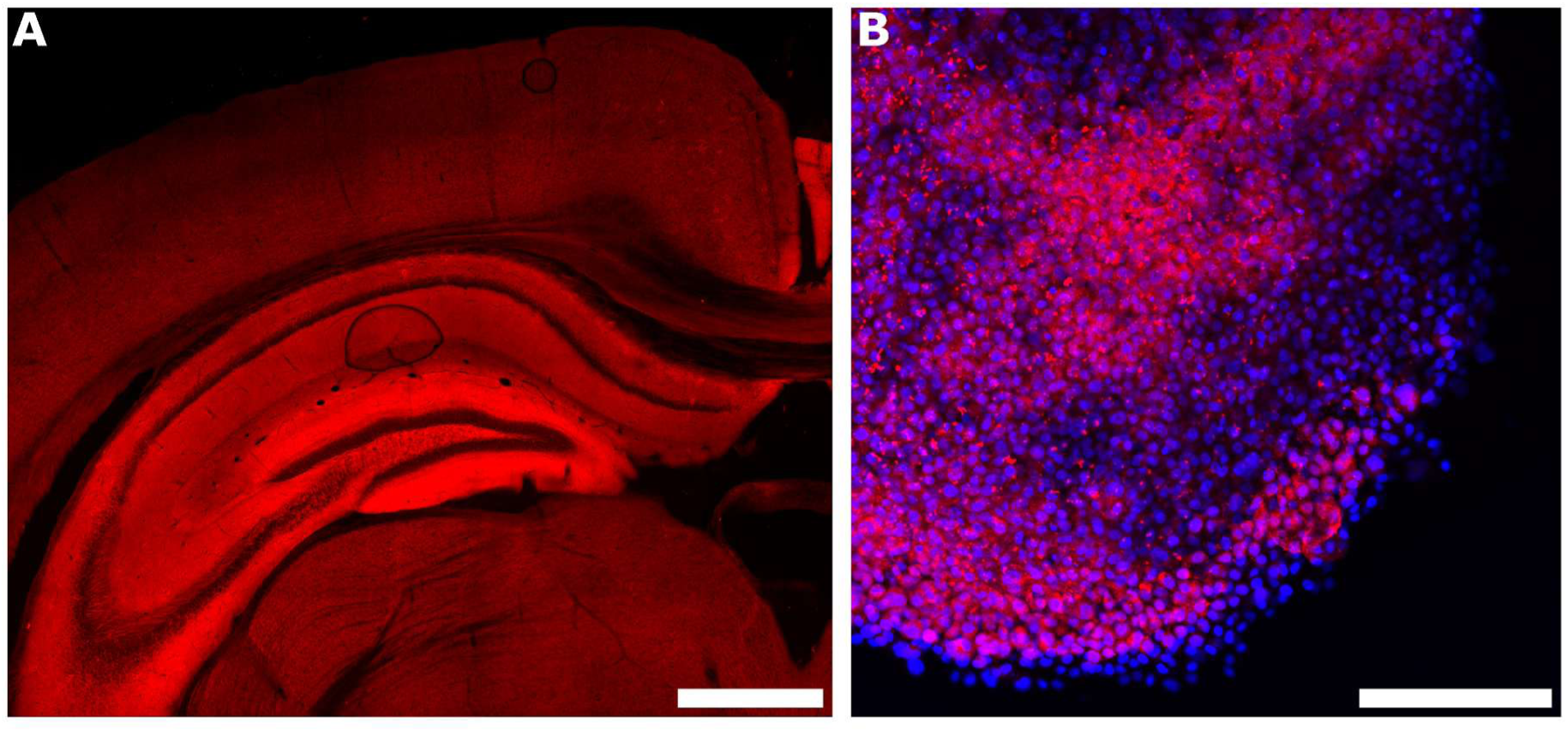
Positive controls for the anti-tenascin-R and anti-tenascin-C antibodies. **(A)** Tenascin-R positive control in hippocampus (dentate gyrus, DG), labeled with an anti-tenascin-R primary antibody and a CF647-conjugated detection antibody (abTenR–abCF647; red); signal is distributed across the hippocampal formation, including the dentate gyrus. **(B)** Tenascin-C positive control in a glioma spheroid, with nuclei counterstained by DAPI (blue) and tenascin-C revealed with an anti-tenascin-C primary antibody and a CF647-conjugated detection antibody (abTenC–abCF647; red). Each antibody yields the expected labeling in tissue in which its target is present, serving as a positive staining control. Scale bars: **A**, 1 mm; **B**, 200 μm.

**Supplementary Figure 3.**
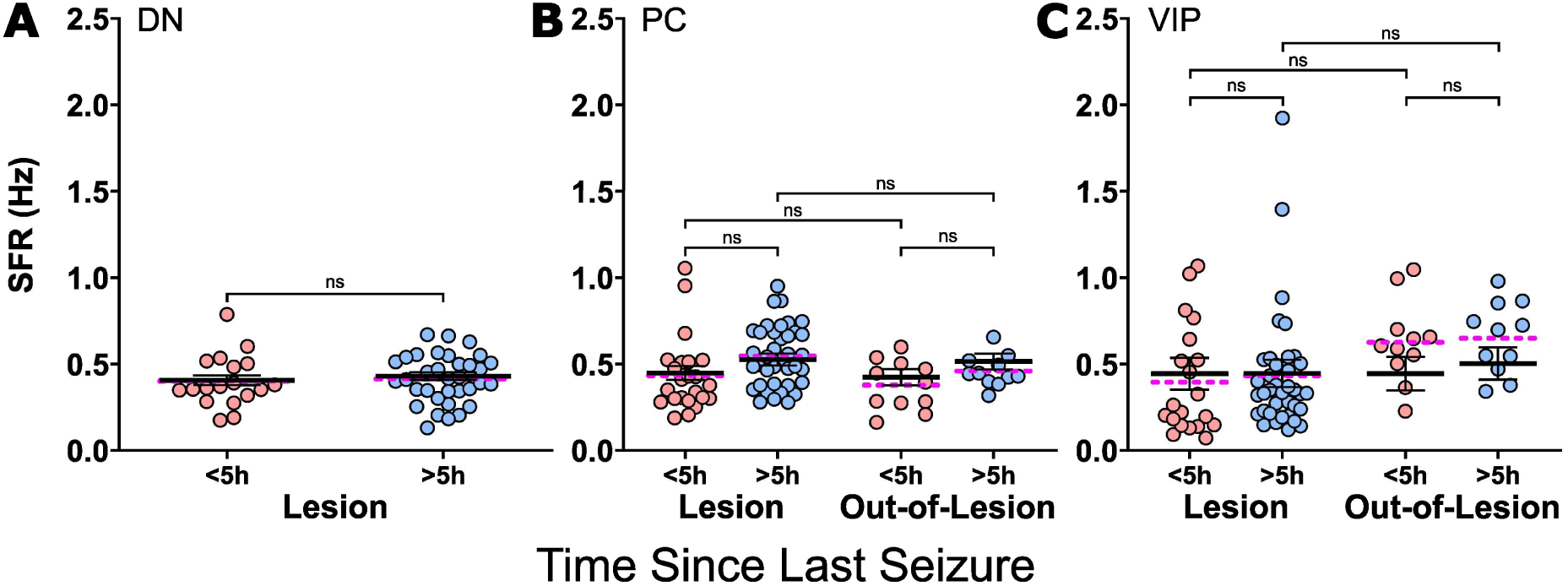
Proximity to the preceding seizure accounts for only a small fraction of the late decline in pyramidal firing. Because SO animals seized only after the early imaging window, their late recordings could fall closer to a seizure than their early ones; this analysis tests whether the decline in Fig. 3G reflects transient post-ictal suppression. Spontaneous firing rate (SFR, Hz) of dysmorphic (DN, **A**), pyramidal (PC, **B**) and VIP (**C**) neurons recorded <5 h (red) or ≥5 h (blue) after the most recent seizure, with the two seizing groups pooled (2 AS, 4 SO). DNs are shown for lesional FOVs only (extra-lesional DN too rare to model); PC and VIP are split into lesional and extra-lesional (Out) FOVs. Each point is one FOV (edge FOVs excluded); solid horizontal bars, model-estimated marginal means; dotted bars, arithmetic FOV means; whiskers, SEM. Brackets denote within-cell-type contrasts, Holm-corrected: the time-window contrast within each location and the lesional-versus-extra-lesional contrast within each window (models in Supplementary Methods). PC: lesional 0.46 ± 0.05 vs 0.54 ± 0.04 Hz (<5 h vs ≥5 h; difference 0.076 ± 0.035 Hz, adjusted *p* = 0.072); extra-lesional 0.41 ± 0.06 vs 0.49 ± 0.06 Hz (0.082 ± 0.052 Hz, adjusted *p* = 0.12); pooled across locations 0.078 ± 0.029 Hz (95% CI 0.020–0.14; *p* = 0.010), about a third of the 0.25 Hz early-to-late decline in Fig. 3G and a little over half of it at the upper 95% bound; no difference between locations in either window (<5 h: 0.049 ± 0.066 Hz; ≥5 h: 0.043 ± 0.064 Hz; adjusted *p* = 0.93 for both). DN: lesional 0.39 ± 0.03 vs 0.41 ± 0.02 Hz (0.021 ± 0.031 Hz, *p* = 0.50). VIP: lesional 0.42 ± 0.12 vs 0.51 ± 0.11 Hz (0.089 ± 0.069 Hz, adjusted *p* = 0.41); extra-lesional 0.37 ± 0.14 vs 0.40 ± 0.14 Hz (0.024 ± 0.096 Hz, *p* = 0.81); no difference between locations (adjusted *p* ≥ 0.55). FOV numbers (<5 h / ≥5 h): DN lesional 18/34; PC lesional 23/38, extra-lesional 11/11; VIP lesional 20/37, extra-lesional 11/11; 4–6 mice per bin.

**Supplementary Figure 4.**
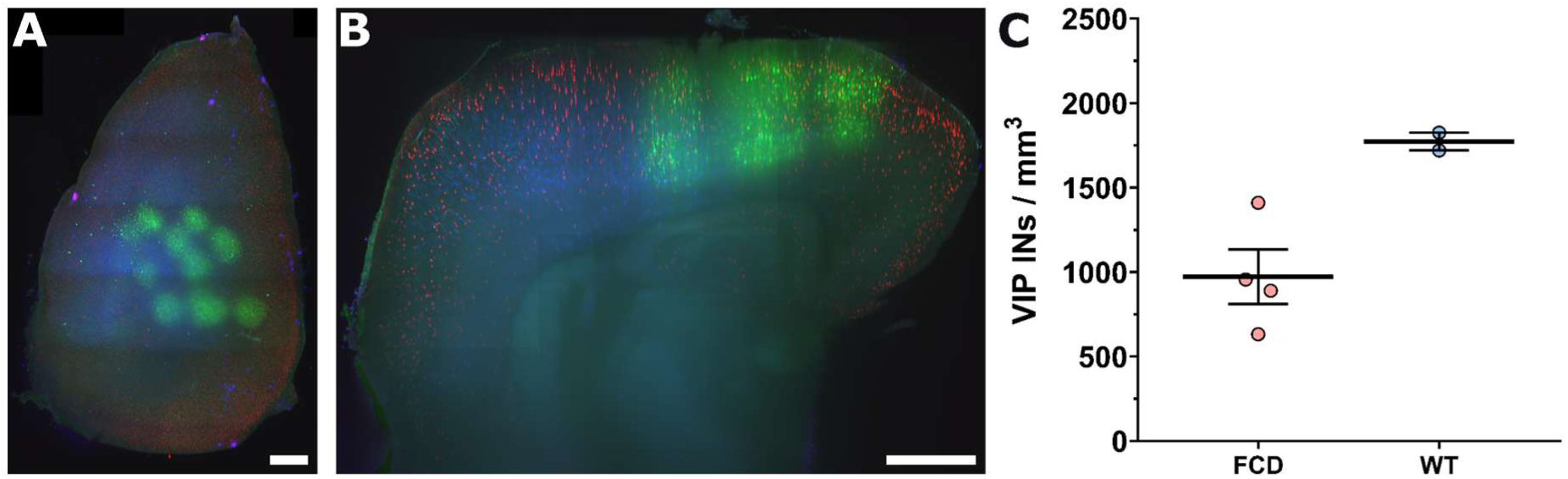
Three-dimensional distribution and spatial density of VIP interneurons in mTOR_WT_ and mTOR_MUT_ brains. (**A**–**B**) Representative light-sheet microscopy dataset from a CUBIC-cleared mTOR-MUT brain, displayed as a maximum-intensity projection from dorsal view **(A)** and a coronal section within the AAV-transduced areas **(B)**. Green, jGCaMP7f-expressing neurons; blue, iRFP_713_-labeled electroporated neurons delineating the FCD lesion; red, tdTomato-positive VIP interneurons. **(C)** Spatial density of tdTomato-positive VIP interneurons (VIP⁺INs/mm³) in mTOR_WT_ control brains (n = 2) and mTOR_MUT_ FCD brains (n = 4). Seizing and non-seizing mTOR_MUT_ animals were pooled for this analysis. VIP-interneuron density was approximately 45% lower in mTOR_MUT_ than in mTOR_WT_ brains. Scale bars: 1 mm; mean ± SEM.

**Supplementary Figure 5.**
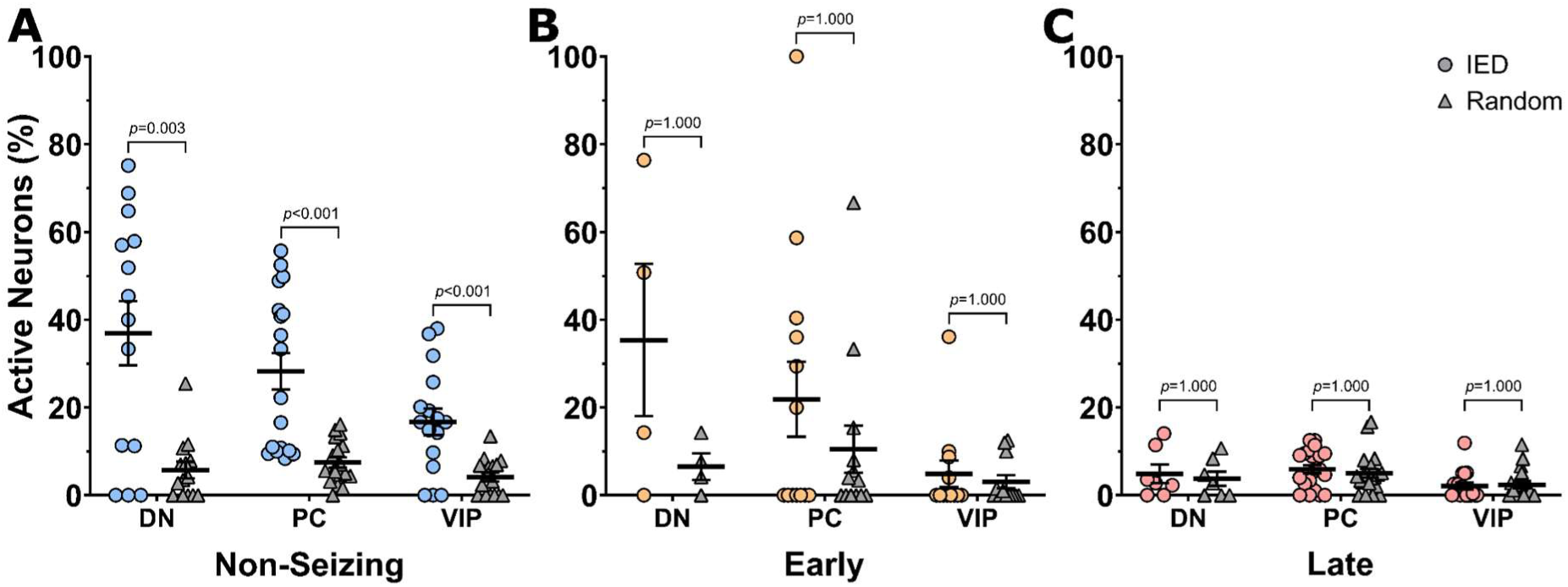
Active-neuron fraction before real versus random IEDs, by condition. Complementary view of the data in Fig. 4F, showing the paired real-versus-random contrast within each disease stage. Fraction of active neurons per field of view (%), defined as neurons firing ≥1 CASCADE-inferred spike in the five frames preceding an event (binary CASCADE output; see Supplementary Methods), for real IEDs (colored circles) versus the same neurons realigned to random times (grey triangles). Panels, condition (non-seizing, early, late seizing tissue); within each panel, points are grouped by cell type—DNs, PCs, and VIP interneurons—and color-coded by condition as in Fig. 4E. Each point is one field of view; horizontal bars, across-FOV means. Brackets compare real-IED versus random-time active fractions for each cell type; *p* values are two-sided paired permutation tests (sign-flip on per-FOV differences, 10⁵ resamples), Holm-corrected across the three cell types within each condition. A significant excess of neurons active before real IEDs is present in non-seizing tissue in all three cell classes (DN *p* = 0.003, PC *p* < 0.001, VIP *p* < 0.001) and is absent in early and late tissue (all *p* > 0.99), consistent with the loss of the pre-IED lead in Fig. 4E and the decline in recruited fraction in Fig. 4F. Per-condition FOV counts differ across cell types; the early-stage DN group in particular is small (n = 4 FOV) and its non-significant result is correspondingly low-powered.

