## Supplementary Methods for "Longitudinal Imaging in a Mouse Model of Focal Cortical Dysplasia Reveals Lesion Expansion and Loss of Pre-Discharge Neuronal Recruitment"

### Sample preparation

#### *Experimental animals*

All experiments were performed under the Animal Care and Animal Protection Law of the Czech Republic, fully compatible with the guidelines of the European Union directive 2010/63/EU. The protocol was approved by the Ethics Committee of the Second Faculty of Medicine (Project License No. MSMT-31765/2019–4). Animals were housed under standard conditions (22 ± 1 °C, 12 h light/dark cycle) with *ad libitum* access to food and water.

Transgenic mice used for the longitudinal functional experiments were generated by crossing VIP-IRES-Cre mice (strain #010908; Jackson Laboratory) with the tdTomato Cre-reporter line Ai9 (lox-STOP-lox-tdTomato; LSL-tdTomato; strain #007909; Jackson Laboratory), resulting in selective expression of the red fluorescent protein tdTomato in VIP interneurons. A separate cohort of five mice on a C57BL/6J background was electroporated with the mutant mTOR construct and used for histological analyses of tenascin-R and tenascin-C (see below). Both male and female mice were included in the study.

A total of 24 mice were included in the study. The longitudinal functional cohort comprised 19 VIP-IRES-Cre × Ai9 mice that underwent the experimental pipeline outlined in Fig. 1C, including *in utero* electroporation, cranial-window implantation, continuous video-EEG monitoring, and repeated two-photon calcium imaging. Of these, 14 mice were electroporated with the mutant mTOR construct (p.L2427P; plasmids provided by SoVarGen Co., Ltd.[1] and further adapted) and five with the wild-type mTOR construct. Among the 14 included mTOR-mutant mice, eight remained free of spontaneous seizures throughout the video-EEG monitoring period and were classified as non-seizing animals (NS; n = 8), whereas six exhibited spontaneous seizures. Four of the six seizing mice developed their first spontaneous seizure between the early and late imaging windows and were classified as seizure-onset animals (SO; n = 4). The remaining two mice were already seizing at the beginning of the longitudinal imaging period and continued to exhibit seizures throughout the experiment (already-seizing animals, AS; n = 2). The five mice electroporated with the wild-type mTOR construct served as mTOR-WT controls (WT; n = 5).

The histological cohort consisted of seizing mTOR-mutant FCD mice and did not contribute to the functional imaging analysis. The mutant mTOR construct was co-electroporated with pCAG-EGFP rather than pCAG-iRFP$_{713}$, allowing direct identification of electroporated neurons in fixed tissue.

#### *In utero electroporation*

*In utero* electroporation (IUE) was performed at embryonic day 14.5 as previously described[2]. Briefly, for animals included in the longitudinal functional experiments, a plasmid mixture consisting of pCAG-iRFP$_{713}$ (2 µg/µL) and either pCAG-mTOR$_{WT}$ or pCAG-mTOR$_{MUT}$ (3 µg/µL) was injected into the left lateral ventricle of each embryo. For the histological cohort, pCAG-iRFP$_{713}$ was replaced with pCAG-EGFP (2 µg/µL), and all embryos were electroporated with pCAG-mTOR$_{MUT}$ (3 µg/µL). All plasmid mixtures were supplemented with 0.03% Fast Green (cat. no. F7252; Sigma-Aldrich).

Electroporation was performed using forceps-type platinum electrodes (CUY650P3, 3 mm; Nepa Gene) connected to a custom-built, current-limited electroporator (current limit, 40 mA; five 35-V pulses of 50 ms duration delivered at 950-ms intervals)[3]. For perioperative analgesia, ketoprofen (5 mg/kg) was administered subcutaneously during surgery and repeated on the first and second postoperative days. Successful electroporation was confirmed by epifluorescence screening at postnatal days 1–2, followed by a second screening at postnatal day 21. Only pups displaying a clearly unilateral fluorescent lesion localized to the somatosensory cortex were retained for further experiments.

***Cranial window implantation***

Mice were operated on at 5–6 weeks of age under isoflurane anesthesia (4–5% induction, 1.5–2% maintenance in 0.5 L/min $O_2$) on a 37 °C heated pad. A ring-shaped titanium headbar with a preassembled connector was affixed to the skull over the left parietal cortex using cyanoacrylate (CA) gel glue (Loctite #1363589; Henkel, Düsseldorf, Germany). Using a handheld high-speed microdrill (Success40; Osada Electric, Tokyo, Japan), we drilled a 4.5 mm craniotomy over the FCD lesion. The bone flap was gently removed to expose the intact dura.

Prior to window placement, AAV was delivered to enable calcium imaging of neuronal activity. A total of 2 × 30 nL of AAV9 (pGP-AAV-syn-jGCaMP7f-WPRE; $5\times10^{11}$ gc/mL) was injected at each of 10 sites into cortical layer 2/3 (250 µm depth) using beveled glass capillaries (3-000-203-G/X; Drummond Scientific, Broomall, USA; 30° angle, 10–15 µm outer diameter) at a rate of 1 nL/s via a Nanoject III injector (Drummond Scientific). The capillary was held in place for 5 min after each injection to allow diffusion and prevent AAV tract leakage. Injection sites were selected to include cortical regions both within and outside the FCD lesion. The craniotomy was then sealed with a 4 mm round #1.5 cover glass (CG15F1, Thorlabs, Newton, USA) placed directly onto the exposed dura and fixed to the surrounding skull with CA glue. We implanted three wire electrodes (51 µm bare / 102 µm coated diameter, PFA insulation; cat. no. 776000, A-M Systems). The lesional electrode was bared over the distal 1 mm, beveled at 45°, and inserted into the parenchyma through the gap between the cranial window and the skull so that its tip lay within the FCD lesion, whereas the contralateral electrode ended in a bare loop (<0.8 mm) resting on the dura over the center of the right parietal bone. A common reference/ground electrode, also ending in a bare loop, was placed under the skull over the cerebellum. For perioperative analgesia, ketoprofen (5 mg/kg) was administered subcutaneously during surgery and repeated on the first and second postoperative days.

***Tissue preparation***

At 17 weeks of age, the animals were deeply anesthetized by an intraperitoneal injection of ketamine (Narketan 10%; 50 mg/kg) and xylazine (Rometar 2%; 6 mg/kg) and transcardially perfused with ice-cold saline followed by 4% paraformaldehyde (PFA) in phosphate-buffered saline (PBS). The brains were removed and post-fixed overnight in 4% PFA at 4 °C. The fixed brains were sectioned coronally at a thickness of 60 µm using a vibratome (Leica VT1200 S; Leica Microsystems, Wetzlar, Germany), and the sections were processed free-floating.

***Immunohistochemistry***

Vibratome sections were processed for immunohistochemistry as described previously[2]. Free-floating sections were washed in PBS, and non-specific binding sites were blocked for 5 h at room temperature using 10% ChemiBLOCKER (Sigma-Aldrich, USA) and 0.5% Triton X-100 in PBS. The sections were subsequently incubated overnight at 4 °C with either a rabbit recombinant monoclonal antibody against tenascin-R (1:150; clone Rb619; cat. no. 217 008; Synaptic Systems) or a rabbit monoclonal antibody against tenascin-C (1:200; clone EPR4219; cat. no. ab108930; Abcam). Following washing in PBS, the sections were incubated overnight at 4 °C with a highly cross-adsorbed CF647-conjugated goat anti-rabbit IgG (H+L) secondary antibody (1:500; cat. no. SAB4600185; Sigma-Aldrich). The sections were then washed in PBS and mounted using Fluoromount-G Mounting Medium (cat. no. 00-4958-02; Thermo Fisher Scientific). Native EGFP fluorescence was used to identify electroporated neurons, and no nuclear counterstain was applied.

#### *Tissue clearing for light-sheet microscopy*

Brains intended for light-sheet microscopy were cleared using the passive-immersion CUBIC method according to Susaki et al.[4]. Following dissection, the brains were post-fixed in 4% PFA at 4 °C for 24 h. The samples were subsequently washed in PBS for at least 24 h, with the buffer renewed every 12 h. Tissue delipidation was performed by incubation in CUBIC-1 solution at room temperature for 1–2 weeks, with fresh solution provided every 3 days. The brains were then extensively washed in PBS for 24 h, replacing the buffer every 3 h. For refractive index matching, the samples were immersed in CUBIC-2 solution for 5 days, with the solution exchanged daily.

#### *Preparation of senescent U-87 MG glioma spheroids*

A glioma spheroid generated from senescence-induced U-87 MG cells (ATCC, Manassas, VA, USA) was used as a positive control for tenascin-C immunostaining (Supplementary Fig. 2). Cellular senescence was induced by treating the cells with 100 µM temozolomide every 3 days for a total of 15 days. For spheroid formation, 50,000–100,000 cells were seeded per well into U-bottom wells of a CELLSTAR® Cell-Repellent 96-well plate (Greiner Bio-One, Frickenhausen, Germany) in 200 µL of Dulbecco's modified Eagle medium containing 4.5 g/L glucose (DMEM; Invitrogen/Gibco, Grand Island, NY, USA), supplemented with 10% fetal bovine serum, 100 U/mL penicillin, and 100 µg/mL streptomycin. The plate was centrifuged at 200 × g for 5 min and subsequently incubated for 3 days at 37 °C in a humidified atmosphere containing 5% $CO_2$ while being agitated at 500 rpm on an MS3 orbital shaker (IKA, Staufen, Germany).

The resulting glioma spheroids were fixed in 4% PFA for 1 h at room temperature and sectioned into 60-µm-thick slices using a vibratome (Leica VT1200 S; Leica Microsystems, Wetzlar, Germany). The sections were processed as free-floating sections and immunostained for tenascin-C using the same protocol as that applied to the brain sections. Briefly, sections were incubated with a rabbit anti-tenascin-C primary antibody (1:200; cat. no. ab108930; Abcam), followed by a goat anti-rabbit CF647-conjugated secondary antibody (1:500; cat. no. SAB4600185; Sigma-Aldrich). After immunostaining, the sections were mounted on microscope slides using a DAPI-containing mounting medium (cat. no. 00-4959-52; Thermo Fisher Scientific).

### Data acquisition

#### *Video-EEG monitoring*

Following a three-week post-surgical recovery period, animals were individually housed in transparent recording chambers and underwent long-term video-EEG monitoring from postnatal week 8–9 until week 17. Monitoring was continuous between weekly two-photon imaging sessions and was conducted under controlled conditions (22 ± 1 °C, 12 h light/dark cycle). EEG was recorded from three platinum/iridium wire electrodes. The imaging field of view (FOV) lay 1–3 mm from the lesional electrode; all analyzed EEG signals were taken from that electrode. Electrophysiological signals were amplified, band-pass filtered at 0.1 Hz–1.6 kHz, and digitized at 5 kHz using a 32-channel headstage with RHD2132 amplifier chip (cat. no. C3314, Intan Technologies, Los Angeles, CA, USA).

The headstage was connected to a custom-built motorized commutator that allowed the animals to move freely within the recording chamber. Vertical movement was accommodated by a self-balancing arm supporting the commutator. Digitized signals were transferred to a PC via an RHD2000 USB Interface Evaluation Board (Intan Technologies) and recorded using Spike2 software (version 10.09b; Cambridge Electronic Design, Cambridge, UK). Behavior was recorded continuously by video alongside the EEG and was used solely to confirm or reject candidate electrographic seizures; no behavioral seizure scoring was performed.

After one week of baseline video-EEG monitoring, animals underwent weekly two-photon calcium imaging under awake head fixation with simultaneous EEG acquisition. During imaging, EEG signals were low-pass filtered at 3 kHz and amplified using a DC-coupled differential amplifier (Model 3000; A-M Systems, Inc., Carlsborg, WA, USA). Signals were digitized at 20 kHz using an NI PXIe-6341 DAQ card in NI PXIe-1073 chassis, connected through a BNC-2090A terminal block (National Instruments, Austin, TX, USA) and recorded in WaveSurfer (version 0.942; Janelia Research Campus, Ashburn, VA, USA). Following each imaging session, animals were allowed one day of recovery before video-EEG monitoring resumed. This alternating monitoring and imaging protocol continued until postnatal week 17. In the recorded EEG signal, seizures were labelled manually using Open Signal Explorer and Labeller (OSEL; https://github.com/kudlaceksystem/OSEL-4_0) written in MATLAB 2019b computing environment (MathWorks; Natick, USA). Seizures were identified by trained experts. They usually exhibited high-frequency pattern at the beginning and transitioned to high-amplitude spiking pattern.

Mice of the histological cohort underwent no surgery and were not implanted with electrodes. Seizures were therefore assessed behaviorally: from postnatal week 12 until perfusion at 17 weeks of age, animals were housed in a custom-built phenotyping cage equipped with piezoelectric sensors that continuously recorded cage vibration alongside synchronized video (manuscript in preparation). An automatic algorithm flagged periods of excessive vibration, and these periods were reviewed on video by a human annotator; the vibration signal served only to select epochs for review, and all seizures were confirmed behaviorally. Events were counted as seizures only when they comprised an apparent loss of posture together with clonic movements. All five animals exhibited such seizures. Note that, unlike the longitudinal cohort, seizure status in this cohort was established behaviorally rather than electrographically.

***Two-photon calcium imaging***

For intravital two-photon calcium imaging, we used a custom-made high-resolution two-photon microscope. The microscope was equipped with a tunable femtosecond laser (Chameleon Vision-S; Coherent, Saxonburg, USA). Laser power and blanking were controlled using an electro-optical modulator (Model 350-80-LA-02 KD*P driven by a 320RM amplifier; Conoptics, Danbury, USA). The angular range of the scanner engine (LSK-GR08; Thorlabs, Newton, USA) was adjusted to produce a square FOV of 810×810 µm at 1× zoom and 405×405 µm at 2× zoom when using a 20× high-NA objective (XLUMPLFLN 20×, NA 1.0, WD 2.0 mm; Olympus, Tokyo, Japan). We recorded neurons from depths ranging from 150 µm to 300 µm below the dura mater.

Fluorescence emission was collected through a 2″ detection pathway, spectrally separated into green and red channels, and detected by two silicon photomultipliers (C13366-3050GA; Hamamatsu, Hamamatsu City, Japan). The signals were preamplified using custom-made video amplifiers (±1 V output) and digitized/recorded using an NI 5732 digitizer connected to an NI PXIe-7961 FPGA and NI PXIe-1073 chassis (National Instruments). Data acquisition, scanning, and laser control were performed using the ScanImage software suite (Vidrio, Ashburn, USA)[5].

Approximately four weeks after surgery, jGCaMP7f expression reached sufficient levels for reliable calcium-imaging recordings. Animals were placed in a custom-made holder and head-fixed via the titanium headbar. A custom-made laser-reflection device was used to precisely level the cranial window surface to the horizontal plane. This step was essential for repeated imaging across longitudinal sessions to ensure relocation of the same neuronal population.

Using a low-magnification objective (TL4X-SAP, 4× Super Apochromatic Objective, NA 0.2; Thorlabs), we acquired an overview image of the entire cortical area under the cranial window, showing the best-transduced regions (920 nm excitation) and their spatial relation to the iRFP-labeled lesion (840 nm excitation). In each mouse we selected 6–10 recording sites, which were then imaged at higher magnification with the 20× objective. To ensure reliable relocation of the same regions across longitudinal sessions, a set of reference images was collected during the first week of imaging and used in all subsequent sessions to identify the same recording sites.

Individual FOVs (405×405 µm at 2× zoom) were classified, relative to the iRFP-delineated lesion border in the overview image, as lying fully inside, at the edge of, or fully outside the lesion, and are termed lesional, edge and extra-lesional ("out"), respectively. In each FOV, two 5-min recordings were acquired weekly. Imaging was performed at <100 mW under the objective, at 30 frames per second and 512×512 pixels, using 920 nm excitation.

EEG was recorded simultaneously with imaging. To synchronize the two acquisition systems, a collateral of the y-galvanometer command signal (sawtooth) was fed to the EEG DAQ as a separate channel; each sawtooth cycle corresponds to one imaging frame, and frame onsets were recovered offline from the flyback transitions. Because both systems sampled the same physical signal, imaging frames and EEG were aligned without drift, to within one frame (33 ms).

#### *Confocal imaging of immunostained sections*

Immunostained sections were imaged on an Andor Dragonfly 503 spinning-disk confocal microscope equipped with a Zyla 4.2 Plus sCMOS camera (Oxford Instruments, UK) and a 20×/0.75 immersion objective (HC PL APO IMM CORR CS2), using identical acquisition settings for all sections and both hemispheres. EGFP was excited at 488 nm (525/50 nm bandpass) and tenascin-C or tenascin-R immunoreactivity at 637 nm (700/75 nm bandpass). Entire coronal sections were imaged, with 20 planes at 1 µm spacing acquired in both channels. Two sections per antibody were imaged in each animal (four sections per animal in total).

#### *Light-sheet microscopy*

In addition to confocal imaging of immunohistochemically processed tissue, brains from a subset of six animals were examined using light-sheet microscopy to obtain three-dimensional spatial information on the extent of the FCD lesions and their associated neuronal projections. This subset included two mTOR-WT control animals and four mTOR-mutant animals with FCD; seizing and non-seizing animals were considered together for the purposes of three-dimensional visualization. Cleared brains were imaged using a LightSheet Z.1 microscope equipped with a 5×/0.1 NA illumination objective, a quad excitation dichroic filter set for 405, 488, 561, and 640 nm excitation (Zeiss, Germany), and a pco.edge 5.5 sCMOS camera (Excelitas, USA). The entire lesion and its associated projections were acquired in three fluorescence channels. jGCaMP7f-expressing neurons were detected using a 488 nm excitation laser and a BP 525/40 nm emission filter, whereas tdTomato fluorescence was recorded using a 561 nm excitation laser and a BP 595/40 nm emission filter. The iRFP signal was excited using the 640 nm laser and detected in the near-infrared emission range using an LP660 emission filter. These channels were used to visualize the distribution and morphology of jGCaMP7f-expressing and iRFP-labeled neurons, as well as the distribution and morphology of tdTomato-positive interneurons. Fluorescence detection and volumetric imaging were performed using a 5×/0.16 NA detection objective (Zeiss, Germany). Individual image tiles were deconvolved and stitched using Huygens Professional software, version 24.04 (Scientific Volume Imaging, the Netherlands). The resulting three-dimensional brain volumes had a voxel size of 1.17×1.17×4.94 µm. Light-sheet volumes were used for three-dimensional visualization and for a descriptive count of tdTomato$^{+}$ VIP interneuron density; no statistical testing was performed on these data.

### Data Processing

Imaging sessions were numbered 1–8 relative to the onset of imaging. In all longitudinal comparisons, EARLY denotes sessions 1–2 and LATE sessions 7–8, approximately six weeks apart. Functional measures (spontaneous firing rate, peri-IED activity) were averaged across the two sessions of each window, whereas structural measures were derived from single sessions: cortical expansion from the averaged motion-registered recording of each FOV in sessions 2 (baseline) and 8, six weeks apart, and soma size from the 60 µm reference z-stacks acquired over the same FOVs in sessions 2 (EARLY) and 8 (LATE). In SO animals the EARLY window preceded the first detected spontaneous seizure and the LATE window followed it; in NS animals no seizures were detected at any time. AS animals were already seizing before the first imaging session and therefore have no pre-seizure EARLY window; for these animals both EARLY and LATE sessions represent the chronic seizing state. Accordingly, AS animals contribute to the lesion-expansion and soma-size comparisons (group S, together with SO), to the time-since-seizure analysis (Supplementary Fig. 3), and to the Late data in the peri-IED analysis; they are excluded from the EARLY-versus-LATE comparisons of firing rate (Fig. 3F–H).

The mean inferred event rate per neuron is referred to as the spontaneous firing rate (SFR); throughout, SFR denotes the activity inferred from jGCaMP7f fluorescence using CASCADE (see Calcium signal processing) rather than directly measured action potentials.

#### *Calcium signal processing*

All two-photon image sequences were motion registered using the NoRMCorre algorithm[6]. The motion-registered movies of each FOV were averaged to obtain a high-quality image from each recorded sequence, which were then used to label neuronal regions of interest (ROI). In the reference FOV image, the ROIs were manually selected in the green channel. Afterwards, we performed polynomial 2D transformation of manually selected key points to map the reference ROIs to the respective FOVs recorded in other imaging sessions.

We categorized the ROIs into 3 types: dysmorphic neurons (DNs), pyramidal cells (PCs), and VIP-positive interneurons (VIPs). Initially, the reference ROIs were manually labeled as VIP-positive or VIP-negative based on fluorescence in the red channel. The red ones were labeled VIPs, while the remaining neurons were sorted according to ROI size. Those larger than 400 pixels (~18 µm in diameter) were marked as DNs and those smaller than 300 pixels (~15 µm in diameter) as PCs. ROIs of intermediate size (300–400 pixels) were not assigned to either class and were excluded from cell-type-specific analyses.

Fluorescence signals were extracted as the mean pixel intensity within each ROI and neuropil signal from surrounding pixels was subtracted using a standard 0.7 coefficient. The baseline of the signal was computed by sequentially filtering the signal with Gaussian, minimum, and maximum filters[7]. Each fluorescence trace was normalized using its baseline signal to obtain relative changes in fluorescence ($(F-F_0)/F_0$). Neuronal spiking activity was inferred from ΔF/F traces using the CASCADE deep learning-based spike inference algorithm[8] using the GCaMP8-tuned model[9]. Among the models available in CASCADE, the GCaMP8-tuned model (with all GCaMP8 variants included) showed highest performance on a GCaMP7f dataset as benchmarked in the later study[9].

Noise level, defined as the median absolute difference between consecutive samples scaled by the square root of the frame rate was calculated for each ΔF/F trace[8]. Only those ΔF/F traces with a noise level lower than 5 %·Hz^(-1/2) were used for further analysis. To assess the spontaneous firing rate, inferred spike rates were averaged across all neurons of the same type within each FOV recording.

To analyze interictal epileptiform discharges (IEDs), we ran the LFP recordings acquired during the imaging through an automatic detector[10] and manually verified all detections to remove false positives. The precise IED time was taken as the peak of maximum voltage deflection. We extracted a time span of 1 second before and after each IED from the imaging data for visualization. Spiking activity traces were averaged in each FOV across all neurons of the same type for all detected IEDs. Imaging was performed at 30 frames per second (one frame ≈ 33 ms), and all peri-IED windows were defined in frames: for each

IED, inferred spiking was summed over the 5 frames immediately preceding the discharge (pre) and the 5 frames immediately following it (post), with the single frame containing the discharge assigned to neither window. Pre-versus-post activity was then compared as described in Statistical analyses.

To analyze the proportion of active neurons, we transformed the inferred spike rates into discrete spikes using the tool included in the CASCADE package[8]. Pre-IED active neurons were defined as those with at least one discrete spike within 5 frames before the IED occurred. Per FOV, the fraction of neurons active in the 167 ms (5 frames) before each event (≥1 discrete CASCADE spike; ≥3 qualifying neurons required) was computed for real IEDs and for the same neurons at matched random times (FOV as unit). In each recording, 60 frames before and after each real IED were removed. From the remaining frames, we randomly selected as many frames as real IEDs in the respective recording. Real-versus-random fractions were then compared as described in *Statistical analyses*.

***Tissue expansion analysis***

For each FOV, the motion-registered functional recording was averaged to obtain a high-quality image of the FOV in the second and eighth imaging sessions, i.e., six weeks apart. These images were then registered using affine transformation with the ITKElastix[11] package in Python 3.11. The resulting transformation matrix was used to calculate scaling factors in x and y axes, which represent tissue expansion in mediolateral (ML) and anteroposterior (AP) axes respectively.

***Single-neuron soma-size analysis***

In a subset of five mice with FCD, we acquired a restricted 60 µm z-stack ranging from +30 µm above to -30 µm below the FOVs used for calcium imaging. Z-stacks encompassing the same set of FOVs were recorded during session 2 (Early) and session 8 (Late) of the imaging protocol. The z-stacks were acquired with a 1 µm z-step, and each image was generated by averaging 60 frames. A custom MATLAB script was used to display the Early and Late stacks for a single FOV side by side on a computer screen, with their left-right positioning randomized. Consequently, the human annotator was blinded to the chronological order (Early vs. Late) of the displayed stacks. A human annotator manually selected the 3–4 largest (presumed DNs) and 3–4 smallest neurons (presumed non-mutated pyramidal cells) that could be identified in both z-stacks. These neurons were then manually delineated in the focal plane where they occupied the largest cross-sectional area. Finally, we calculated the effective diameter as the diameter of a circle with an equivalent area.

***Tenascin quantification***

Confocal stacks of immunostained sections (see Confocal imaging of immunostained sections) were collapsed into maximum-intensity projections and analyzed in MATLAB R2024a. The lesion ROI was defined on the EGFP projection by Gaussian blurring ($\sigma = 3$ px) followed by an automatic Otsu threshold, delineating the cortical area occupied by electroporated neurons. For normalization, a homotopic, cortex-restricted region of equal area in the contralateral hemisphere was segmented manually, blind to the tenascin channel. Mean gray values of the tenascin projection were measured in both ROIs; no background subtraction was applied, as any additive offset affects both measurements equally. Tenascin signal is reported as the lesion-to-contralateral mean gray value ratio. Ratios from the two sections stained for the same antibody were averaged, yielding one tenascin-C and one tenascin-R value per animal (n = 5).

#### *Statistical analyses*

Linear mixed-effects models were fitted in MATLAB R2024a (Statistics and Machine Learning Toolbox 24.1) using fitlme with restricted maximum likelihood; fixed-effect contrasts were tested with Satterthwaite-approximated denominator degrees of freedom, and omnibus effects by likelihood-ratio tests on maximum-likelihood fits. Other tests were performed in GraphPad Prism (v10.4), MATLAB R2024a or Python 3.11.5, with permutation procedures implemented as custom scripts. All tests were two-sided. Descriptive values are mean ± s.e.m. (or median [IQR] where noted); model-derived values are estimated marginal means (EMMs) ± s.e.m. Families of related comparisons were Holm-corrected within the family defined for each analysis below.

Unless stated otherwise, the unit of analysis is the FOV. Each FOV contributes one value to each analyzed condition — imaging window, group or time-since-seizure bin — obtained by averaging across the neurons of the given cell type and across the recordings assigned to that condition. Numbers of animals and FOVs entering each analysis are given in the respective figure legend; they are smaller than the cohort totals because an animal contributed only if it provided at least one qualifying FOV in all compared conditions.

Significance is denoted throughout all figures by *$p < 0.05$, **$p < 0.01$, ***$p < 0.001$ and ns, not significant; figure legends do not restate this.

Cortical expansion (Fig. 2C). Per-FOV expansion relative to session 2 was computed along the mediolateral (ML) and anteroposterior (AP) axes for WT (n = 6 FOVs), seizing (S = AS + SO; n = 22) and NS (n = 22) animals. FOVs were included when both sessions were imaged and the affine registration converged. Only lesional FOVs were considered. In WT animals, which develop no lesion, the corresponding FOVs lay within the iRFP-labeled electroporated area. A single family of twelve comparisons was Holm-corrected: per group and axis, a one-sample t-test versus 0% (annotated on the x-axis); and, per axis, all three between-group pairs by Welch's t-test (effect sizes, Hedges' g).

Tenascin (Fig. 2F). Per animal, the lesion-to-contralateral intensity ratio (one value per animal, two sections averaged; n = 5) was tested against unity by a two-sided one-sample t-test for each protein (tenascin-C, tenascin-R), Holm-corrected across the two.

Somatic growth (Fig. 2I). Effective soma diameter (see Single-neuron soma-size analysis) was compared between sessions 2 and 8 for DN (n = 30 FOVs) and PC (n = 26) by two-sided paired t-tests with the FOV as the unit, Holm-corrected across the two. To test whether the two cell classes differed in growth, the per-FOV difference in growth (ΔDN − ΔPC) was computed in the 26 FOVs containing both classes and tested against zero by a two-sided one-sample t-test; this contrast was not included in the Holm family above.

SFR — cross-sectional (Fig. 3D, E). SFR was analyzed with the FOV as the unit (per-FOV mean per cell type; EARLY window; FCD animals only, SO and NS pooled, 12 animals; lesional and extra-lesional FOVs; edge FOVs excluded). DN and PC (Fig. 3D; groups DN-lesion n = 41, PC-lesion n = 50, PC-out n = 35) were modeled as SFR ~ cell + (1|mouse) + (1|FOV) (REML), with an overall likelihood-ratio test (2 d.f.) and three pre-specified pairwise contrasts (Holm). VIP cells (Fig. 3E; one value per FOV; lesion n = 50, out n = 35) were modeled as SFR ~ location + (1|mouse), without a FOV intercept, and tested by a likelihood-ratio test (1 d.f.).

SFR — longitudinal (Fig. 3F–H). This analysis was restricted to lesional FOVs. A separate model was fitted per neuron type and group (PC and VIP: WT, SO, NS; DN: SO and NS only; AS animals and edge FOVs excluded): SFR ~ time point + (1|mouse) + (1|FOV) (REML), with the FOV as the unit. Because not every FOV contributed a qualifying recording in both windows, the numbers of EARLY and LATE observations differ; the FOV random intercept accommodates this incomplete pairing. The EARLY-versus-LATE contrast was Holm-corrected across groups within each panel.

Firing versus time since seizure (Supplementary Fig. 3). In seizing animals (AS + SO; 6 animals), recordings were binned by the interval since the most recent seizure into near (<5 h) and distant (≥5 h) windows; the 5 h boundary was chosen a priori. SFR was modeled with the FOV as the unit and with animal and FOV random intercepts: SFR ~ window × location + (1|mouse) + (1|FOV) for PC and VIP, and SFR ~ window + (1|mouse) + (1|FOV) for DN (lesional FOVs only). Within each cell type, two contrast families (window within location; location within window) were Holm-corrected. Because the two locations yielded nearly identical window shifts in PC, a reduced model without the location term (SFR ~ window + (1|mouse) + (1|FOV)) was also fitted for PC to estimate the pooled window effect; this single contrast was pre-specified and reported uncorrected.

Peri-IED activity (Fig. 4E). Per IED and neuron, d = pre − post (summed over the five frames after minus the five frames before the discharge; windows defined in Data Processing). With the FOV as the unit of replication, the per-FOV mean d was tested against zero by a two-sided one-sample t-test for each cell type and condition. Conditions were: non-seizing (NS animals), early (SO animals before their first spontaneous seizure; session 1+2) and late (SO animals after their first spontaneous seizure, together with AS animals; session 7+8). Holm correction was applied across cell types within each condition, in two families: lesional FOVs (DN, PC, VIP) and extra-lesional FOVs (PC, VIP).

Active-neuron fraction (Fig. 4F, Supplementary Fig. 5). For each lesional FOV, the fraction of neurons active in the five frames (~167 ms) preceding each event (≥1 discrete CASCADE spike; at least three qualifying neurons of the given type required) was computed for real IEDs and for matched random times (see Data Processing), with the FOV as the unit. Supplementary Fig. 5: real-versus-random fractions were compared within each cell type and condition by a two-sided paired permutation test (sign-flip on per-FOV differences, $10^5$ resamples), Holm-corrected across the three cell types within each condition. Fig. 4F: the real-IED fraction was compared between conditions by a two-sided permutation test (label shuffling, $10^5$ resamples), Holm-corrected across the three pairwise condition comparisons within each cell type. Because the sets of qualifying FOVs differed between conditions, these between-condition comparisons were treated as independent rather than paired.

Descriptive data. Seizure frequency and mean seizure duration (Supplementary Fig. 1) and VIP interneuron density (Supplementary Fig. 4C) are reported descriptively and were not tested statistically; no inferential claim is made from these values.

**SUPPLEMENTARY METHODS - references**
